# E-cigarette vapour induces cellular senescence in primary lung fibroblasts and may contribute to lung pathology

**DOI:** 10.1101/2023.08.30.555286

**Authors:** Jack Bozier, Baoming Wang, Maaike de Vries, Corry-Anke Brandsma, Maarten van den Berge, Wim Timens, Irene H. Heijink, Brian G.G. Oliver, Roy R. Woldhuis

## Abstract

Cellular senescence has been recognized to play a role in COPD pathophysiology. Non-aerosolized E-liquid treatment of a lung fibroblast foetal cell line resulted in cellular senescence induction, but this has not been assessed using E-vapour exposure and using primary lung fibroblasts. Therefore, we investigated whether E-vapour exposure induces cellular senescence in primary human lung fibroblasts and whether this affects their tissue repair function.

Primary human lung fibroblasts were stimulated with E-cigarette vapour extracts, and cigarette smoke extract (CSE) and Paraquat (PQ) as positive controls. IL-8 secretion was measured to confirm a stimulatory response. Multiple senescence markers (p16, p21, SA-β-gal, and proliferation) were assessed and the tissue repair function was assessed using a scratch assay. Finally, we tried to validate our findings in an E-cigarette-exposed mouse model.

Upon stimulation with CSE, PQ, and E-vapour extracts, cellular senescence was induced, which seemed dose-dependent and nicotine-independent. E-vapour-induced senescence resulted in an impaired tissue repair function. No significant difference was observed in the E-cigarette-exposed mouse model.

In this study, we identify E-cigarette vapours’ potential to induce cellular senescence in primary human lung fibroblasts and that this affects their tissue repair function, which further adds to the identified risks of E-cigarette use.

## INTRODUCTION

COPD is a progressive inflammatory lung disease caused by exposure to noxious gases, in particular cigarette smoke. Chronic exposure to cigarette smoke (CS) causes inflammation, leading to lung tissue damage. Ultimately, these pathologic events lead to airway fibrosis and airflow limitation, as well as alveolar breakdown and lack of tissue repair in the parenchyma, i.e., emphysema.(1)

Cellular senescence has been recognized to play a role in the pathophysiology of COPD.(2) Cellular senescence is defined as an irreversible cell cycle arrest to prevent cell death or abnormal growth. Induction of cellular senescence can be caused by multiple mechanisms, including oxidative stress and DNA damage,(3) both known to result from chronic CS exposure.(4) CS has been demonstrated to induce cellular senescence in multiple structural lung cells,(5, 6) including lung fibroblasts,(7, 8) and *in vivo* in murine lungs and human lungs.(4, 9) Accumulation of senescent cells in lung tissue can result in chronic inflammation and tissue dysfunction,(3) thus contributing to COPD pathology.(2)

Electronic cigarettes have been proposed as a safer alternative to cigarettes. However, evidence of the harms related to E-cigarette use is growing. Similar to CS, E-cigarette exposure causes a pro-inflammatory response after acute and chronic exposure in multiple structural lung cells *in vitro, in vivo* in mouse lungs, and in human clinical studies,(10) suggesting prolonged use may also contribute to COPD. Furthermore, E-cigarette vapour has been shown to induce DNA damage and reduce DNA damage repair in lung epithelial cell lines.(11) Oxidative stress from E-cigarette exposure has been studied to a greater extent *in vitro* and *in vivo*, demonstrating a high oxidative burden from E-vapour exposure.(12) Although E-cigarette use may up-regulate DNA damage and oxidative stress in epithelial and fibroblast cell lines, it is unknown whether this is the case in primary human lung fibroblasts. Furthermore, studies investigating cellular senescence due to E-cigarette exposure have been performed in a foetal cell line utilizing non-aerosolized E-liquids, which is less representative of exposure conditions in the lung.(13-15) To the best of our knowledge no studies have been performed to assess cellular senescence upon E-vapour exposure in primary human lung fibroblasts. Therefore, we investigated whether E-vapour exposure induces cellular senescence in primary human lung fibroblasts and whether this affects their tissue repair function.

## METHODS

### Primary parenchymal lung fibroblast isolation

Primary lung fibroblasts from subjects undergoing lung transplantation or tumour resection surgery were used. Primary parenchymal lung fibroblasts were isolated and cultured from parenchymal lung tissue as described previously.(16) Briefly, parenchymal lung tissue was cut into small cubes and cultured in tissue flasks in DMEM medium supplemented with 10% foetal bovine serum (FBS), and 2% antibiotics at 37°C and 5% CO_2_. Medium was refreshed every week and when cultures reached confluency, fibroblasts were trypsinized and passaged into new culture flasks or frozen and stored in liquid nitrogen.

### E-vapour stimulation

Primary human parenchymal lung fibroblasts from non-COPD and COPD patients (n=8) were grown in DMEM (Gibco) supplemented with 5% FBS and 2% antibiotics at 37°C and 5% CO_2_. Cells of passage 5-6 were seeded on plates for 48 hours prior to serum starvation in 0.5% FBS DMEM, and 24 hours later cells were stimulated with 250 μM Paraquat (Sigma-Aldrich) (PQ; positive control for senescence induction), 5% cigarette smoke extract (CSE), 1.5% (Lo) or 2% (Hi) nicotine-containing (18mg/ml) tobacco-flavoured E-cigarette Vapour extract (EV), or 1.5% (Lo) or 2% (Hi) nicotine-free tobacco-flavoured E-cigarette Vapour extract (NF EV) as described previously.(17) These concentrations of CSE, EV, and NF EV were not cytotoxic (data not shown). A cytotoxic dose for CSE (10%) and both EV (5%) were used as positive controls for the stimuli (data not shown). Supernatants (for IL-8 ELISA, BD Biosciences) and RNA extracts were collected after 24 hours of stimulation, whilst the remaining plates were refreshed to 5% FBS DMEM to enable cell proliferation for 3 days for SA-β-gal staining and wound healing assay.

### Cellular senescence assays

Cellular senescence was assessed by SA-β-gal staining, cell proliferation inhibition, and p16 and p21 gene expression as described previously.(18) Briefly, cells were fixed with 2% paraformaldehyde + 0.2% Glutaraldehyde for 5 minutes at room temperature. After washing with PBS, cells were stained with SA-β-gal staining solution for 16 hours at 37°C without CO_2_. The next day, cells were stained with 1ug/ml DAPI and four representative areas per well were imaged on a Nikon Eclipse Ti microscope at 100x total magnification. Total cells (DAPI positive nuclei) and SA-β-gal positive cells were manually counted to calculate percentages of SA-β-gal positive cells.

For p16 and p21 gene expression, RNA was isolated using Isolate II RNA Mini Kit (Bioline) and cDNA was synthesized using SensiFAST cDNA Synthesis kit (Bioline). Gene expression was measured using qRT-PCR using TaqMan (Applied Biosystems) assays and SensiFAST™ Probe Hi-ROX Kit (Bioline) according to the manufacturer’s protocol. Relative expression was calculated by 2(-ΔCp) with 18S as reference gene.

### Wound healing assay

Wound healing capacity was assessed 4 days after exposure using a scratch assay. A wound was made in the confluent fibroblast layer by scratching with a p200 pipet tip in the middle of the well from top to bottom. After scratching, the wells were washed twice with Hank’s buffer, and DMEM + 0.5% FBS was added to enable wound closure. The wound area was measured 48 and 72 hours after scratching, which was captured on a Nikon Eclipse Ti microscope at a total magnification of 40x. Wound healing was calculated by measuring the size of closure corrected for the size of the initial wound, which was expressed as a percentage of closure.

### Validation of E-vapour-induced senescence *in vivo*

To validate E-vapour-induced senescence *in vivo*, samples of a previously described E-vapour exposure model were used.(19) RNA was isolated from lung homogenates from mice exposed to E-cigarettes (with and without nicotine) and room air (SHAM) using Isolate II RNA Mini Kit (Bioline) and cDNA was synthesized using SensiFAST cDNA Synthesis kit (Bioline). Gene expression was measured using qRT-PCR using TaqMan (Applied Biosystems) assays and SensiFAST™ Probe Hi-ROX Kit (Bioline) according to the manufacturer’s protocol. Relative expression was calculated by 2(-ΔCp) with 18S as reference gene.

### Statistical analyses

All stimulated lung fibroblast cultures were compared to untreated lung fibroblast cultures using Friedman tests with Dunn’s post-hoc tests. Samples from E-vapour-exposed mice were compared to room air-exposed mice using a Kruskal-Wallis test with Dunn’s post-hoc tests. P-value < 0.05 was considered statistically significant.

## RESULTS

### Stimulatory response of CSE and E-vapour on primary lung fibroblasts

As cigarette smoke extract (CSE) and E-vapour extracts (EV) are variable and unstable stimuli, we first confirmed the stimulatory response of the different stimuli on primary lung fibroblasts by measuring IL-8 secretion. Il-8 secretion was significantly induced by almost all stimuli compared to untreated fibroblasts (Figure 1). Only the low dose of nicotine-free E-vapour extract resulted in a trend (p=0.058) toward a significant increase (Figure 1B).

**Figure 1:**
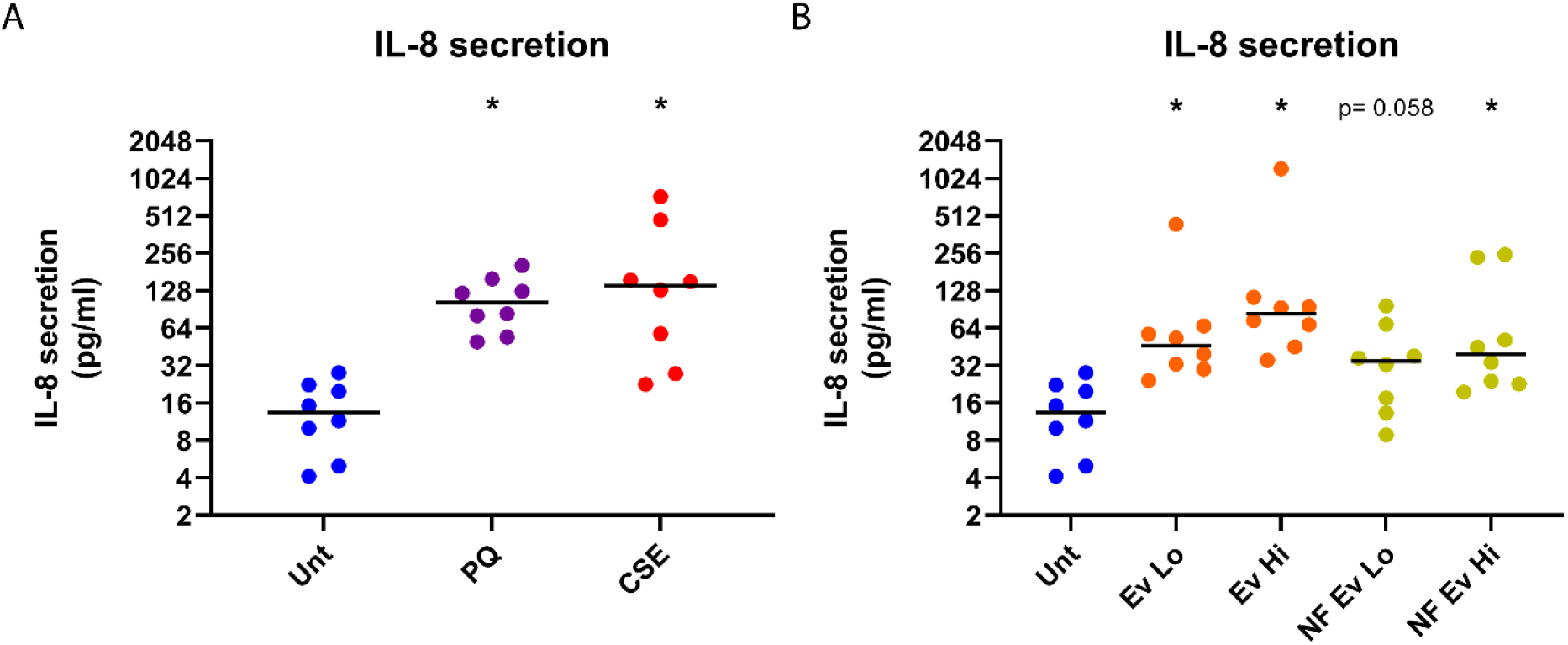
Stimulatory response of CSE and E-vapour on primary lung fibroblasts. IL-8 protein secretion 24h post-stimulation of known-senescence inducers (A) and E-vapour extracts (B) are depicted in scatter plots. Untreated fibroblasts (Unt) are depicted in blue and fibroblasts stimulated with 250μM paraquat (PQ, purple) 5% cigarette smoke extract (CSE, red), and low (Lo = 1.5%) and high (Hi = 2%) doses of nicotine-containing (EV Lo and Hi, orange) and nicotine-free E-vapour (NF EV Lo and Hi, yellow) extracts are depicted. Lines represent medians. N=8 per group. Significant differences between stimulated and untreated fibroblasts were tested using Friedman tests with Dunn’s post-hoc tests. * means p-value < 0.05 compared to untreated. Trends towards significance (p = 0.05 – 0.1) are indicated with p-values.

### E-vapour-induced cellular senescence in primary lung fibroblasts

Paraquat (PQ) and CSE were used as positive controls for senescence induction. Indeed, cellular senescence was induced by PQ and CSE with increased p21 expression (Fig. 2A) and percentages of SA-β-gal positive cells (Fig. 2B), and reduced cell proliferation (Fig. 2C). For both stimulations, no increase in p16 expression was found (data not shown). CS is a known risk factor for COPD and both stimuli are known to induce senescence, confirming our model’s validity.

**Figure 2:**
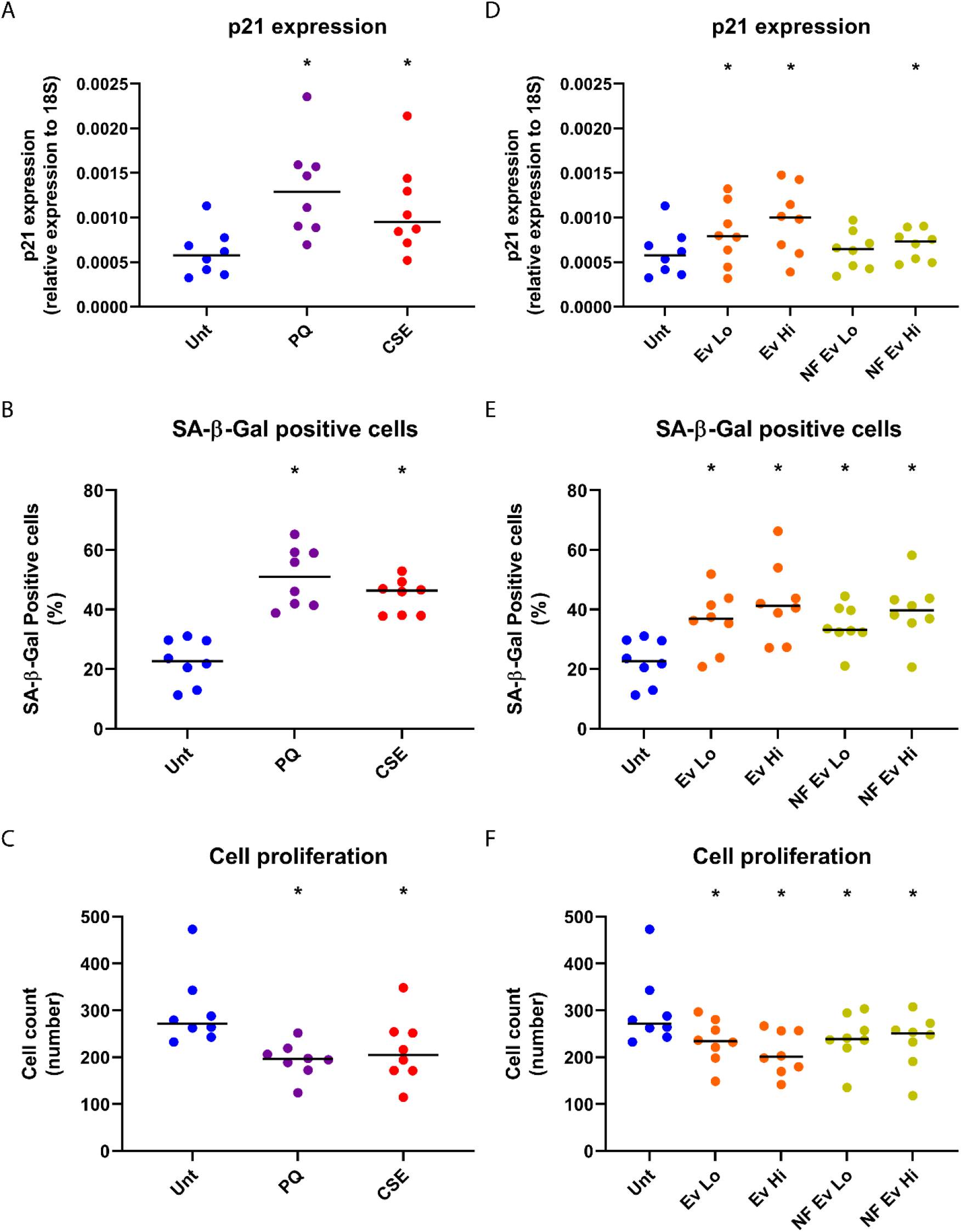
E-vapour-induced cellular senescence in primary lung fibroblasts. Senescence markers after stimulation with known-senescence inducers (A-C) and E-vapour extracts (D-F) are depicted in scatter plots. p21 gene expression (A+D) (24h post-stimulation), percentages of Senescence-associated beta-galactosidase (SA-β-gal) positive cells (B+E) and total cell numbers (C+F) (4d post-stimulation) are depicted. Untreated fibroblasts (Unt) are shown in blue and fibroblasts stimulated with 250μM paraquat (PQ, purple) 5% cigarette smoke extract (CSE, red), and low (Lo = 1.5%) and high (Hi = 2%) doses of nicotine-containing (EV Lo and Hi, orange) and nicotine-free E-vapour (NF EV Lo and Hi, yellow) extracts are depicted. Lines represent medians. N = 8 per group. Significant differences between stimulated and untreated fibroblasts were tested using Friedman tests with Dunn’s post-hoc tests. * means p-value < 0.05 compared to untreated.

Upon stimulation with nicotine-containing E-vapour extract, cellular senescence was induced with significant differences in the same senescence markers (p21, SA-β-gal & proliferation) as induced by PQ and CSE, which seemed dose-dependent (Fig. 2D-F). Nicotine-free E-vapour extract (NF EV) also induced senescence, as shown by increased p21 expression and percentages of SA-β-gal positive cells, along with reduced cell proliferation. Only upon the low dose of NF EV stimulation, no significant p21 increase was observed (Fig. 2D). So, the fibroblasts seem to be slightly more sensitive to senescence induction by nicotine-containing E-vapour extract compared to E-vapour extract without nicotine. Similar to PQ and CSE, no increase in p16 expression was found after NF EV and EV stimulations (data not shown).

### E-vapour-induced senescence resulted in impaired tissue repair function of primary lung fibroblasts

To assess the effect of E-vapour-induced senescence on the tissue repair function of primary lung fibroblasts we used a scratch wound healing assay. Upon senescence induction by PQ and CSE, impaired tissue repair in the wound healing model occurred after 48 hours (Fig. 3A) and 72 hours, with only a trend (p=0.058) for CSE (Fig. 3B). Both doses of EV impaired wound healing capacity with reduced wound closure after 48 hours (Fig. 3C) or 72 hours (Fig. 3D), which appeared to be nicotine-independent as comparable results were found in NF EV stimulated cells, with both doses having significant effects at both time-points (Fig. 3C+D). No significant differences were observed for Ev Lo after 48h and for Ev Hi after 72h, due to lower group size caused by technical issues.

**Figure 3:**
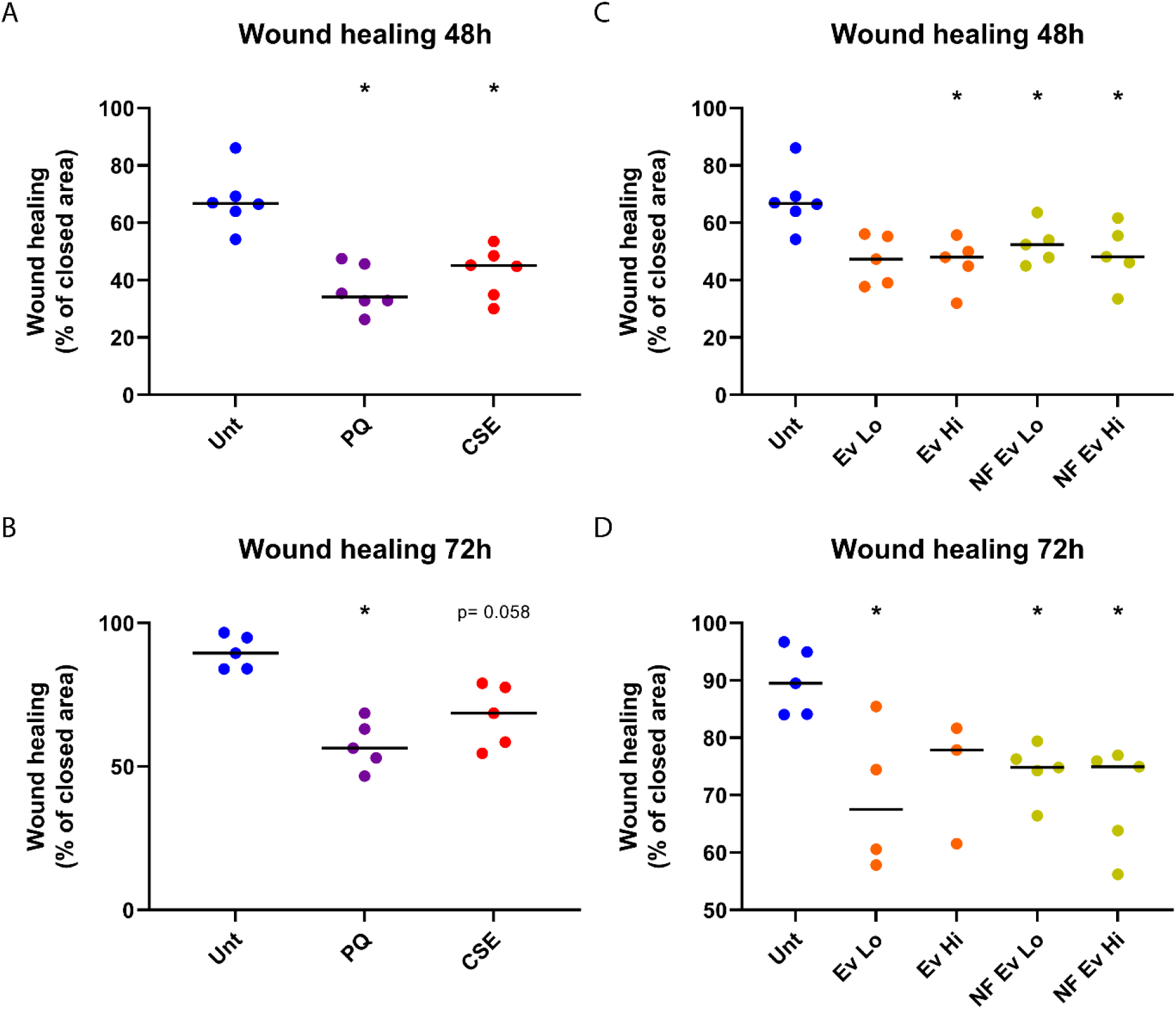
E-vapour-induced senescence resulted in impaired tissue repair function of primary lung fibroblasts. Wound healing after stimulation with known-senescence inducers (A+B) and E-vapour extracts (C+D) after 48 hours (A+C) and 72 hours (B+D) are depicted in scatter plots. Untreated fibroblasts (Unt) are depicted in blue and fibroblasts stimulated with 250μM paraquat (PQ, purple) 5% cigarette smoke extract (CSE, red), and low (Lo = 1.5%) and high (Hi = 2%) doses of nicotine-containing (EV Lo and Hi, orange) and nicotine-free E-vapour (NF EV Lo and Hi, yellow) extracts are depicted. Lines represent medians. Technical outliers, including scratch failures, were excluded. N=3-6 per group. Significant differences between stimulated and untreated fibroblasts were tested using Friedman tests with Dunn’s post-hoc tests. * means p-value < 0.05 compared to untreated. Trends towards significance (p = 0.05 – 0.1) are indicated with p-values.

### Validation of E-vapour-induced senescence *in vivo*

To confirm our *in vitro* results *in vivo*, gene expression of p21 was measured in a previously performed E-cigarette-exposed mouse model,(19) where p21 expression appeared slightly higher upon exposure compared to non-exposed controls (Figure 4). However, no significant differences were found due to high variance and because this pilot study was underpowered, thus larger *in vivo* studies should be done to confirm our *in vitro* findings.

**Figure 4:**
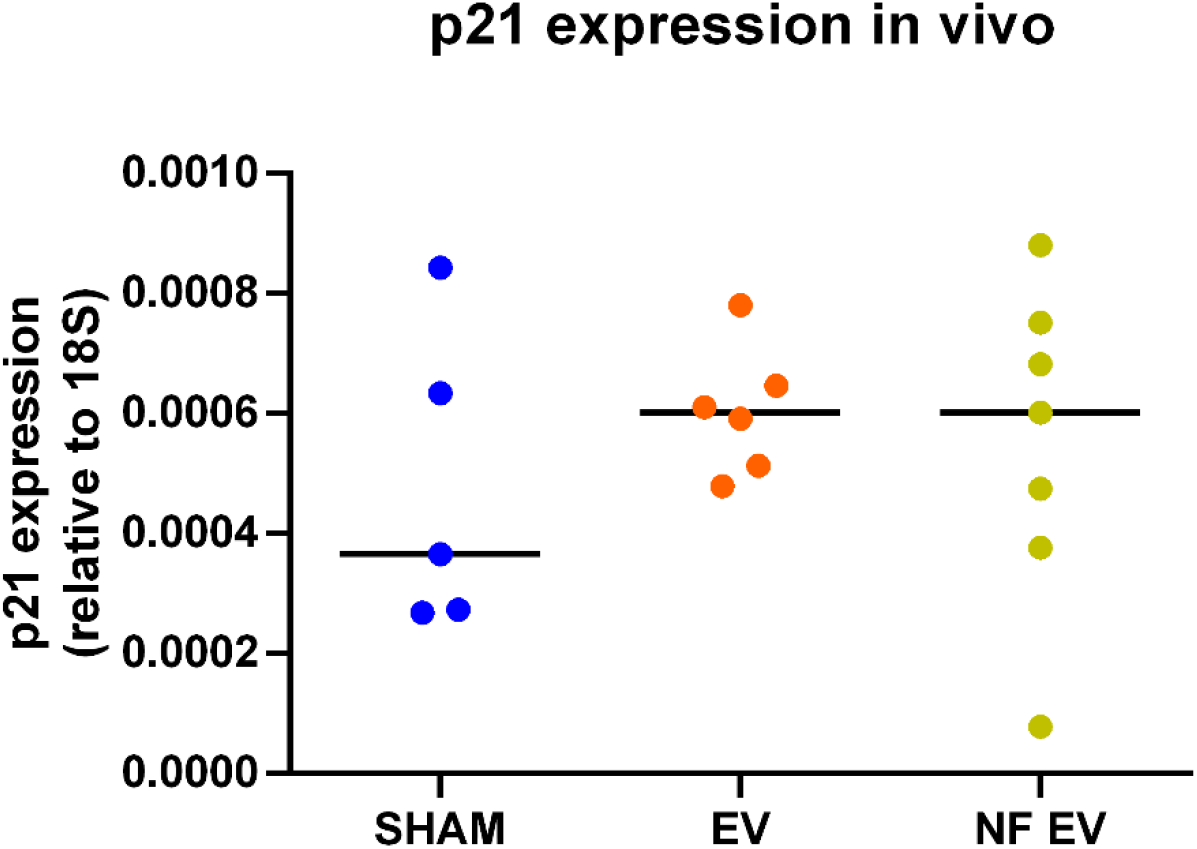
P21 gene expression upon E-vapour exposure in vivo in mice. Relative p21 gene expression of room air-exposed mice (SHAM, blue) and mice exposed with nicotine-containing E-vapour (EV, orange) and nicotine-free E-vapour (NF EV, yellow) is depicted. Lines represent medians. N=5-7 per group. Significant differences were tested using a Kruskal–Wallis test.

## DISCUSSION

This study identifies E-cigarette vapours’ potential to induce cellular senescence in primary human lung fibroblasts, which is a known contributing factor to disease in COPD.(2) Previous studies have reported E-cigarette-induced senescence in non-lung cells or human lung (foetal) cell lines, but not in primary lung fibroblasts and these studies did not attempt to replicate the variety of ways in which e-cigarettes are used (high and low doses, and with and without nicotine). The findings of our study further add to the identified risks of short-term E-cigarette use.(10) E-cigarette harms are often compared to cigarettes in relation to harm reduction, but this study focused on the standalone risk for E-cigarette users. Chronic long-term use of tobacco cigarettes is well-known to cause lung tissue damage including cellular senescence. Our results should be a warning that long-term E-cigarette use can also contribute to the induction of cellular senescence in primary lung fibroblasts and impact their tissue repair function, which may lead to COPD development. These risks are not isolated to COPD patients, as other E-cigarette users, including young never-smokers, may be more likely to develop lung pathology from long-term use.

In the current study, we did not directly investigate the mechanisms of senescence induction by E-vapour extract, but we hypothesize that DNA damage and oxidative stress may be involved as previous studies demonstrated that E-cigarette vapour exposure can result in DNA damage and oxidative stress in epithelial cells *in vitro*.(10-12, 14) Future studies should elucidate the mechanisms involved in EV-induced senescence and whether specific components of E-liquids are directly up-regulating these mechanisms.

Lung fibroblasts play an important role in lung tissue repair by their supporting role of producing extracellular matrix proteins, matrix metalloproteinases, and growth factors .(1, 20-22) Cellular senescence has been described to have a multifaceted role in tissue repair, firstly in support of wound repair(23-25) and stem cell regeneration,(26) while conversely contributing to a delayed or impaired wound healing process(27-30) and inhibition of stem cell regeneration.(31) In mice models, p16-positive senescent murine lung fibroblasts promote epithelial stem cell regeneration after injury,(32) while a recent study demonstrated that senescence-induced human lung fibroblasts impaired murine alveolar epithelial cell progenitor function.(33) These studies suggest that there might be a tight balance, where transient senescence is supportive for tissue repair and persistent senescence is detrimental for tissue repair.(34) This balance may be tissue and (micro-) environment-dependent. In this study, we demonstrated that E-vapour-induced senescent human lung fibroblasts have an impaired tissue repair function. Impaired tissue repair has been recognized to be a feature of COPD pathology.(1, 21, 35) Whether E-vapour-induced senescence-impaired tissue repair contributes to COPD pathology *in vivo* is unknown and should be elucidated using more complex 3D *ex vivo* models, like precision-cut lung slices.

E-cigarettes’ potential to induce cellular senescence in primary human lung fibroblasts, alongside other previously identified risks, should serve as a warning to avoid use as a safe alternative to cigarette smoking or as a cessation device. Considering senescence induction seemed dose-dependent indicates excessive and long-term use should be avoided.

## List of abbreviations

COPD: Chronic Obstructive Pulmonary Disease
CS: Cigarette Smoke
CSE: Cigarette Smoke Extract
DMEM: Dulbecco’s Modified Eagle Medium
EV: E-vapour extract (nicotine-containing)
FBS: Foetal Bovine Serum
Hi: High concentration
Lo: Low concentration
NF EV: Nicotine-Free E-vapour extract
PQ: Paraquat
SA-β-gal: Senescence-associated beta-galactosidase

## Authors’ contributions

Conceptualization: JB, BGGO, RRW

Methodology: JB, BW, BGGO, RRW

Software: NA

Validation: NA

Formal Analysis: JB, MdV, CAB, MvdB, WT, IHH, BGGO, RRW

Investigation: JB, BW, RRW

Resources: BGGO

Data Curation: JB, BGGO, RRW

Writing – Original Draft Preparation: JB, RRW

Writing – Review & Editing: All authors

Visualization: JB, RRW

Supervision: BGGO

Project Administration: JB, BGGO, RRW

Funding Acquisition: BGGO

RRW and BGGO contributed equally.

## Ethics approval and consent to participate

Ethical approval for all experiments with primary lung cells was provided by the Human Ethics Committees (IRB) of the Royal Prince Alfred Hospital, Sydney and the University of Technology, Sydney and written informed consent was obtained.

## Data availability

The datasets during and/or analysed during the current study are available from the corresponding author upon reasonable request.

## Acknowledgements

We would like to thank Dikaia Xenaki (*Woolcock Institute of Medical Research*) for her contribution to the biobank of primary lung fibroblasts from lung tissue from COPD patients and non-COPD subjects.

## REFERENCES

1. Hogg JC, and Timens W. The pathology of chronic obstructive pulmonary disease. Annu Rev Pathol 4: 435–459, 2009.

2. Barnes PJ, Baker J, and Donnelly LE. Cellular Senescence as a Mechanism and Target in Chronic Lung Diseases. Am J Respir Crit Care Med 200: 556–564, 2019.

3. Muñoz-Espín D, and Serrano M. Cellular senescence: from physiology to pathology. Nat Rev Mol Cell Biol 15: 482–496, 2014.

4. Birch J, Anderson RK, Correia-Melo C, Jurk D, Hewitt G, Marques FM, Green NJ, Moisey E, Birrell MA, Belvisi MG, Black F, Taylor JJ, Fisher AJ, De Soyza A, and Passos JF. DNA damage response at telomeres contributes to lung aging and chronic obstructive pulmonary disease. Am J Physiol Lung Cell Mol Physiol 309: L1124–1137, 2015.

5. Tsuji T, Aoshiba K, and Nagai A. Cigarette smoke induces senescence in alveolar epithelial cells. Am J Respir Cell Mol Biol 31: 643–649, 2004.

6. Ahmad T, Sundar IK, Lerner CA, Gerloff J, Tormos AM, Yao H, and Rahman I. Impaired mitophagy leads to cigarette smoke stress-induced cellular senescence: implications for chronic obstructive pulmonary disease. Faseb j 29: 2912–2929, 2015.

7. Nyunoya T, Monick MM, Klingelhutz A, Yarovinsky TO, Cagley JR, and Hunninghake GW. Cigarette smoke induces cellular senescence. Am J Respir Cell Mol Biol 35: 681–688, 2006.

8. Kanaji N, Basma H, Nelson A, Farid M, Sato T, Nakanishi M, Wang X, Michalski J, Li Y, Gunji Y, Feghali-Bostwick C, Liu X, and Rennard SI. Fibroblasts that resist cigarette smoke-induced senescence acquire profibrotic phenotypes. Am J Physiol Lung Cell Mol Physiol 307: L364–373, 2014.

9. Aoshiba K, Zhou F, Tsuji T, and Nagai A. DNA damage as a molecular link in the pathogenesis of COPD in smokers. Eur Respir J 39: 1368–1376, 2012.

10. Bozier J, Chivers EK, Chapman DG, Larcombe AN, Bastian NA, Masso-Silva JA, Byun MK, McDonald CF, Crotty Alexander LE, and Ween MP. The Evolving Landscape of e-Cigarettes: A Systematic Review of Recent Evidence. Chest 157: 1362–1390, 2020.

11. Ganapathy V, Manyanga J, Brame L, McGuire D, Sadhasivam B, Floyd E, Rubenstein DA, Ramachandran I, Wagener T, and Queimado L. Electronic cigarette aerosols suppress cellular antioxidant defenses and induce significant oxidative DNA damage. PLoS One 12: e0177780, 2017.

12. Zhao J, Zhang Y, Sisler JD, Shaffer J, Leonard SS, Morris AM, Qian Y, Bello D, and Demokritou P. Assessment of reactive oxygen species generated by electronic cigarettes using acellular and cellular approaches. J Hazard Mater 344: 549–557, 2018.

13. Lucas JH, Muthumalage T, Wang Q, Friedman MR, Friedman AE, and Rahman I. E-Liquid Containing a Mixture of Coconut, Vanilla, and Cookie Flavors Causes Cellular Senescence and Dysregulated Repair in Pulmonary Fibroblasts: Implications on Premature Aging. Front Physiol 11: 2020.

14. Lerner CA, Rutagarama P, Ahmad T, Sundar IK, Elder A, and Rahman I. Electronic cigarette aerosols and copper nanoparticles induce mitochondrial stress and promote DNA fragmentation in lung fibroblasts. Biochem Biophys Res Commun 477: 620–625, 2016.

15. Shivalingappa PC, Hole R, Westphal CV, and Vij N. Airway Exposure to E-Cigarette Vapors Impairs Autophagy and Induces Aggresome Formation. Antioxid Redox Signal 24: 186–204, 2016.

16. Krimmer DI, Burgess JK, Wooi TK, Black JL, and Oliver BG. Matrix proteins from smoke-exposed fibroblasts are pro-proliferative. Am J Respir Cell Mol Biol 46: 34–39, 2012.

17. Bozier J, Rutting S, Xenaki D, Peters M, Adcock I, and Oliver BG. Heightened response to e-cigarettes in COPD. ERJ Open Res 5: 2019.

18. Woldhuis RR, de Vries M, Timens W, van den Berge M, Demaria M, Oliver BGG, Heijink IH, and Brandsma CA. Link between increased cellular senescence and extracellular matrix changes in COPD. Am J Physiol Lung Cell Mol Physiol 319: L48–l60, 2020.

19. Chen H, Li G, Chan YL, Chapman DG, Sukjamnong S, Nguyen T, Annissa T, McGrath KC, Sharma P, and Oliver BG. Maternal E-Cigarette Exposure in Mice Alters DNA Methylation and Lung Cytokine Expression in Offspring. Am J Respir Cell Mol Biol 58: 366–377, 2018.

20. Ito Y, Correll K, Schiel JA, Finigan JH, Prekeris R, and Mason RJ. Lung fibroblasts accelerate wound closure in human alveolar epithelial cells through hepatocyte growth factor/c-Met signaling. Am J Physiol Lung Cell Mol Physiol 307: L94–105, 2014.

21. Togo S, Holz O, Liu X, Sugiura H, Kamio K, Wang X, Kawasaki S, Ahn Y, Fredriksson K, Skold CM, Mueller KC, Branscheid D, Welker L, Watz H, Magnussen H, and Rennard SI. Lung fibroblast repair functions in patients with chronic obstructive pulmonary disease are altered by multiple mechanisms. Am J Respir Crit Care Med 178: 248–260, 2008.

22. White ES. Lung extracellular matrix and fibroblast function. Ann Am Thorac Soc 12 Suppl 1: S30–33, 2015.

23. Demaria M, Ohtani N, Youssef SA, Rodier F, Toussaint W, Mitchell JR, Laberge RM, Vijg J, Van Steeg H, Dollé ME, Hoeijmakers JH, de Bruin A, Hara E, and Campisi J. An essential role for senescent cells in optimal wound healing through secretion of PDGF-AA. Dev Cell 31: 722–733, 2014.

24. Jun JI, and Lau LF. The matricellular protein CCN1 induces fibroblast senescence and restricts fibrosis in cutaneous wound healing. Nat Cell Biol 12: 676–685, 2010.

25. Jiang D, de Vries JC, Muschhammer J, Schatz S, Ye H, Hein T, Fidan M, Romanov VS, Rinkevich Y, and Scharffetter-Kochanek K. Local and transient inhibition of p21 expression ameliorates age-related delayed wound healing. Wound Repair Regen 28: 49–60, 2020.

26. Chiche A, Le Roux I, von Joest M, Sakai H, Aguín SB, Cazin C, Salam R, Fiette L, Alegria O, Flamant P, Tajbakhsh S, and Li H. Injury-Induced Senescence Enables In Vivo Reprogramming in Skeletal Muscle. Cell Stem Cell 20: 407–414.e404, 2017.

27. Samdavid Thanapaul RJR, Shvedova M, Shin GH, Crouch J, and Roh DS. Elevated skin senescence in young mice causes delayed wound healing. Geroscience 44: 1871–1878, 2022.

28. Kita A, Saito Y, Miura N, Miyajima M, Yamamoto S, Sato T, Yotsuyanagi T, Fujimiya M, and Chikenji TS. Altered regulation of mesenchymal cell senescence in adipose tissue promotes pathological changes associated with diabetic wound healing. Commun Biol 5: 310, 2022.

29. Wilkinson HN, Clowes C, Banyard KL, Matteuci P, Mace KA, and Hardman MJ. Elevated Local Senescence in Diabetic Wound Healing Is Linked to Pathological Repair via CXCR2. J Invest Dermatol 139: 1171–1181.e1176, 2019.

30. Zhou F, Onizawa S, Nagai A, and Aoshiba K. Epithelial cell senescence impairs repair process and exacerbates inflammation after airway injury. Respir Res 12: 78, 2011.

31. Banito A, Rashid ST, Acosta JC, Li S, Pereira CF, Geti I, Pinho S, Silva JC, Azuara V, Walsh M, Vallier L, and Gil J. Senescence impairs successful reprogramming to pluripotent stem cells. Genes Dev 23: 2134–2139, 2009.

32. Reyes NS, Krasilnikov M, Allen NC, Lee JY, Hyams B, Zhou M, Ravishankar S, Cassandras M, Wang C, Khan I, Matatia P, Johmura Y, Molofsky A, Matthay M, Nakanishi M, Sheppard D, Campisi J, and Peng T. Sentinel p16(INK4a+) cells in the basement membrane form a reparative niche in the lung. Science 378: 192–201, 2022.

33. Bramey N, Melo-Narvaez MC, See F, Ballester-Lllobell B, Steinchen C, Jain E, Hafner K, Yildirim AÖ, Königshoff M, and Lehmann M. Primary human lung fibroblasts exhibit trigger-but not disease-specific cellular senescence and impair alveolar epithelial cell progenitor function. bioRxiv 2023.2007.2024.550385, 2023.

34. Wilkinson HN, and Hardman MJ. Senescence in Wound Repair: Emerging Strategies to Target Chronic Healing Wounds. Front Cell Dev Biol 8: 773, 2020.

35. Chung KF, and Adcock IM. Multifaceted mechanisms in COPD: inflammation, immunity, and tissue repair and destruction. Eur Respir J 31: 1334–1356, 2008.

